# Incorporation of a nucleoside analog maps genome repair sites in post-mitotic human neurons

**DOI:** 10.1101/2020.03.25.008490

**Authors:** Dylan A. Reid, Patrick J. Reed, Johannes C.M. Schlachetzki, Grace Chou, Sahaana Chandran, Ake T. Lu, Claire A. McClain, Jean H. Ooi, Jeffrey R. Jones, Sara B. Linker, Enoch C. Tsui, Anthony S. Ricciardulli, Shong Lau, Simon T. Schafer, Steve Horvath, Jesse R. Dixon, Nasun Hah, Christopher K. Glass, Fred H. Gage

**Author notes:** These authors contributed equally to this work (Dylan A. Reid, Patrick J. Reed). These authors contributed equally to this work (Johannes C.M. Schlachetzki, Grace Chou, Sahaana Chandran).

## Abstract

Neurons are the longest-living cells in our bodies, becoming post-mitotic in early development upon terminal differentiation. Their lack of DNA replication makes them reliant on DNA repair mechanisms to maintain genome fidelity. These repair mechanisms decline with age, potentially giving rise to genomic dysfunction that may influence cognitive decline and neurodegenerative diseases. Despite this challenge, our knowledge of how genome instability emerges and what mechanisms neurons and other long-lived cells may have evolved to protect their genome integrity over the human life span is limited. Using a targeted sequencing approach, we demonstrate that neurons consolidate much of their DNA repair efforts into well-defined hotspots that protect genes that are essential for their identity and function. Our findings provide a basis to understand genome integrity as it relates to aging and disease in the nervous system.

**One Sentence Summary:** Recurrent DNA repair hotspots in neurons are linked to genes essential for identity and function.

## Main Text

Neurons are highly specialized post-mitotic cells comprising the major functional cell type of the central nervous system. While there is a limited capacity to generate new neurons throughout life, the majority of neurons age in parallel with the organism, making them especially susceptible to decline from age-related disruptions in cellular homeostasis (*1, 2*). Neurons must repair ~10^4-5^ DNA lesions each day, amounting to more than one billion over the lifespan (*3*). Deficiencies in DNA repair pathways have been linked to developmental neurodegenerative disorders (*4*) and to genome instability, a primary hallmark of aging often associated with age-related neurodegenerative diseases (*4–7*).

Early work on genome integrity in neurons suggested that DNA repair was primarily focused on transcribed genes at the expense of inactive regions of the genome not essential for neuronal function (*8, 9*). Accumulation of DNA lesions drives age-associated changes in transcription that lead to a decline in neuronal function (*10, 11*). Additionally, neuronal activity correlates with the generation of DNA double strand breaks (DSBs), potentially contributing to genomic instability (*12, 13*). Despite a clear link between genome maintenance and neuronal health, we know surprisingly little about how neurons maintain genome integrity, as most of our knowledge comes from studies of mitotic neural progenitor cells or expensive whole genome sequencing of single neurons to address somatic mosaicism (*14–18*). Most genomic approaches require a substantial number of cells for targeted specific DNA lesion detection, limiting their adoption by the field (*14, 19*). These technical limitations hamper our ability to define the genome protection strategies that neurons have evolved to ensure their unique longevity.

To better understand genome integrity in neurons, we developed a sequencing method capable of capturing the genomic locations of all DNA repair based on the incorporation of the click chemistry nucleoside analog, EdU (5-ethynyl-2’-deoxyuridine). Previous reports have described the ability of neurons to incorporate radioactive thymidine into their genomes following DNA damage or under normal resting conditions by DNA repair pathways (*20, 21*). We confirmed this to be the case by using single-molecule, localization-based, super-resolution imaging of EdU in human embryonic stem cell-induced neurons (ESC-iNs) incubated with EdU for 24 hrs (Fig. 1A & fig. S1A-B) (*22, 23*). These neurons have EdU clusters in both the nucleus and cytosol, where EdU is incorporated into mitochondrial nucleoids during mitochondrial biogenesis. To define the genomic locations of EdU molecules that had been incorporated into the nuclear genome of ~500,000 ESC-iNs fed EdU for 24 hrs, we opted to enrich next-generation sequencing libraries for fragments that contain EdU via click chemistry addition of a biotin epitope (Supplementary Methods). This strategy is similar to the targeted sequencing of newly synthesized DNA containing nucleoside analogs to identify the locations of replication forks (*24, 25*). Our method, termed “Repair-Seq,” revealed many locations across the neuronal genome that exhibited substantially more EdU enrichment over comparable whole-genome sequencing to the same depth (Fig. 1B & fig. S2A-B).

**Fig. 1.**
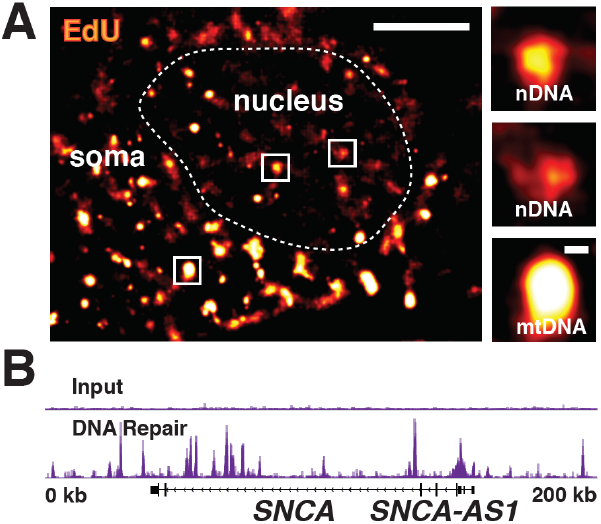
EdU incorporated into the genomes of post-mitotic neurons by DNA repair can be mapped by next-generation sequencing. (A) Representative super-resolution image of ESC-iN nucleus (dashed line) with EdU repair foci and select zoomed regions. Small EdU clusters are evident in the nucleus (nDNA), whereas mitochondrial biogenesis leads to bright mitochondrial nucleoids (mtDNA) in the cell body. (B) DNA repair peaks from the *SNCA* locus in EdU-fed ESC-iNs compared with input genomes sequenced to the same depth show substantial enrichment at some sites. Scale bars are 5 microns and 250 nm respectively.

EdU enriched sites appear as well-defined peaks, so we applied a genome peak calling algorithm to our data, finding ~87,000 total peaks in both H1 and H9 ESC-iNs. We found 61,178 peaks in common for both lines, covering ~1.6% of the genome (fig. S2C-D) (Table S1). As these sites exhibited relative enrichment for DNA repair over the rest of the genome, we termed them DNA repair hotspots (DRHs). These DRHs were distributed throughout the genome on all chromosomes, and appeared to be enriched in promoters, 5’UTRs, and gene bodies (fig. S2E-F). To exhibit such a stable signal in our assay, recurrence across lines and replicates suggests that these locations are frequently repaired in the sequenced ESC-iN population. Additionally, our approach is not specific for any particular DNA repair pathways, instead capturing a heterogenous mix of all repair pathways that are capable of nucleotide incorporation.

Given the stability and reproducibility of these DRHs, we next sought to define the genomic and epigenomic features that could contribute to their establishment in neurons. To map the locations of open chromatin and active regulatory regions in our ESC-iNs, we performed ATAC-Seq and H3K27Ac ChIP-Seq, respectively, and found that ~23.5% of Repair-Seq common peaks were located within these genomic regions (Fig. 2A & fig. S3A-B) (*26*). This percentage to a ~15-fold enrichment over expected associations for repair and these chromatin marks.

**Fig. 2.**
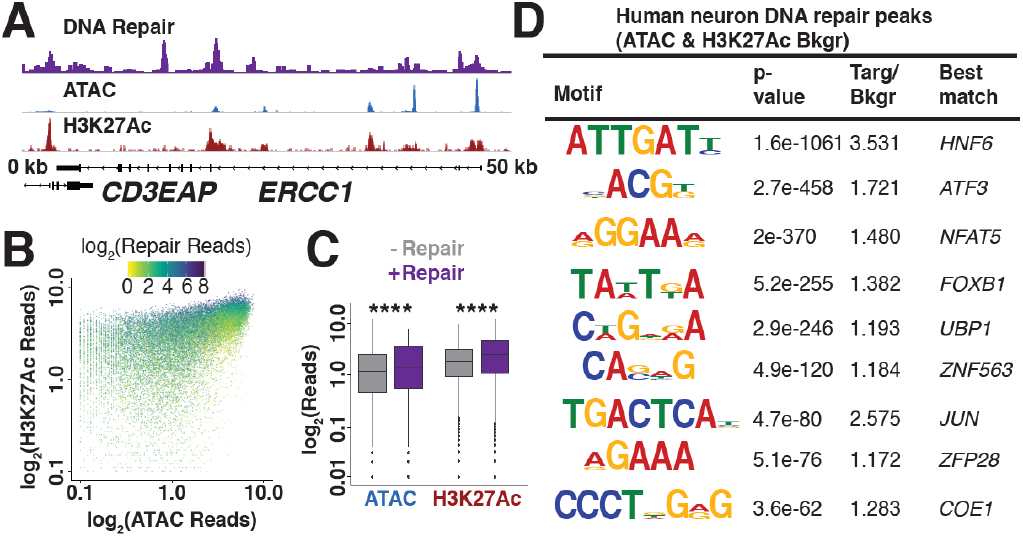
Chromatin accessibility controls the placement of DNA repair hotspots. (A) Repair-, ATAC-, and H3K27Ac ChIP-Seq data at the *ERCC1* locus demonstrate overlap between DNA repair, chromatin accessibility, and histone acetylation. (B) Scatter plot of Repair-Seq normalized read counts compared to ATAC and H3K27Ac normalized read counts. (C) Box plots of ATAC and H3K27Ac peaks with and without DNA repair. (D) DNA sequence motifs identified *de novo* and predicted as enriched in DRHs relative to randomized sequence. **** p-value <2.2e-16 Kruskal-Wallis test.

Intersecting peaks in open regions correlate with greater DNA repair signal strength (Fig. 2B) (*27, 28*). This conclusion is supported by evidence that ATAC and H3K27Ac sites that intersect with DRHs have more normalized reads than those lacking repair (Fig. 2C). Additionally, when we used Repair-Seq peaks as a reference and plot ATAC and H3K27Ac signal intensity, we found that both of these marks, if not directly overlapping, were proximal to DRHs (fig. S3C). Promoters were the predominant point of intersection for Repair, ATAC, and H3K27Ac peaks, whereas DRHs that did not associate with open chromatin were predominantly located in intergenic and intronic elements of the genome (fig. S3D).

To examine the underlying contribution of the genome to the formation of DRHs, we performed *de novo* DNA sequence motif analysis for all peaks. We identified DNA sequence motifs, including ones associated with the factors *HNF6*, *ATF3*, *NFAT5*, *FOXB1*, *UBP1*, *ZNF563*, *JUN*, *ZFP28*, and *COE1* as being significantly enriched in Repair-Seq peaks when taking ATAC and H3K27Ac peaks as background to correct for the contributions of open chromatin (Fig. 2D & fig. S3E-F & Table S2). Many of the factors associated with these motifs have roles in specifying neuronal characteristics (*29–31*). We next asked if our *de novo* DNA repair-associated motifs were enriched in genes with DRHs, compared to genes that did not form DHRs, and we found that these motifs were not enriched in DRH containing genes (fig. S3G). This lack of enrichment suggests that the establishment of DRHs could occur in other genes and that there might be as yet undefined “organizing factors” that coordinate sites of recurrent DNA repair in non-dividing cells.

Repair-Seq allowed us to directly compare all DNA repair- and transcription-associated reads. A majority of Repair-Seq reads (~67%) could be assigned to gene bodies using RNA-Seq pipelines (*32*), with most of the neuronal transcriptome exhibiting some level of maintenance that increased with expression (Fig. 3A & fig. S4A-B). This finding is largely in agreement with prior work suggesting that, in neurons, global DNA repair is attenuated and consolidated to actively transcribed genes, presumably to suppress the accumulation of lesions and mutations (*8*). However, when we examined the reads that comprise DRHs (~23% of all repair reads), we observed that many more genes lacked these recurrent DNA repair sites and we found no relationship to expression (Fig. 3B & fig. S4C) (Table S3). Almost a third of DRHs were located in intergenic regions; therefore, we could not readily correlate these with transcription of single genes. To address the potential contribution of these sites to transcription-associated repair, we generated Hi-C contact maps for ESC-iNs such that we could assign intergenic peaks to genes based on features of 3D genome organization, such as Topologically Associating Domains (TADs) (*33*). Total DNA repair levels in most TADs were uniform (Fig. 3C). Assignment of intergenic peaks did not substantially alter the interpretation that DRHs were not correlated with the levels of gene transcription (Fig. 3D & fig. S6). Comparing the distribution of either all DNA repair-associated reads or Repair-Seq peaks with genome-wide features of 3D genome organization such as A/B compartments, we found an enrichment of DNA repair in the “active” A compartment (fig. S5A-C).

**Fig. 3.**
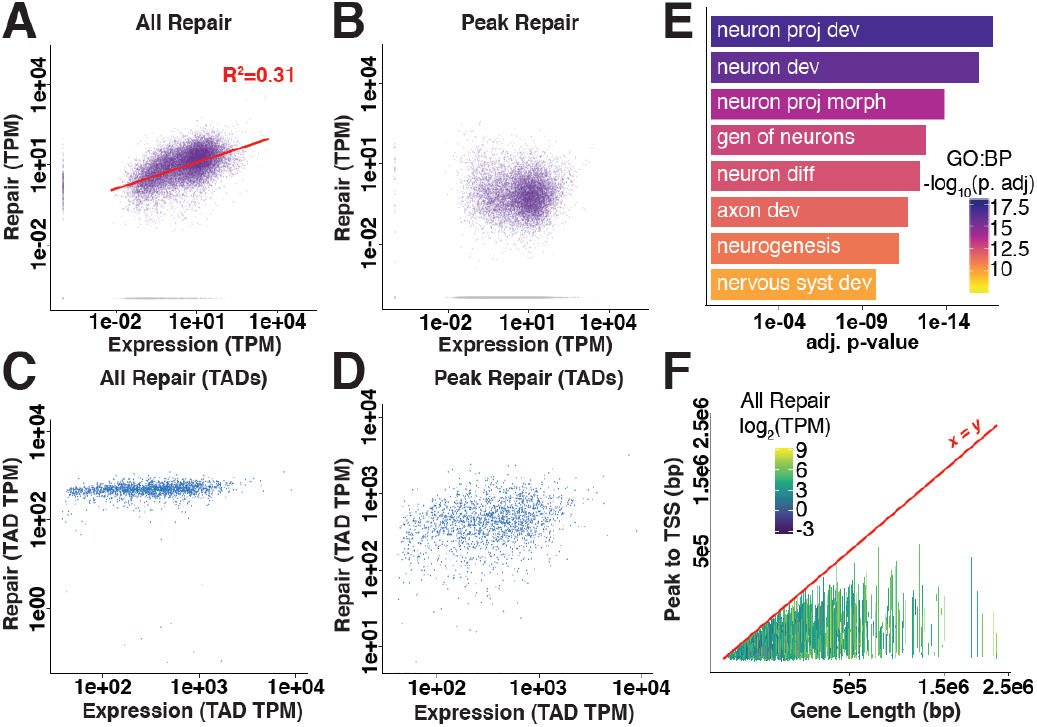
Transcriptional output correlates with total DNA repair in genes but not DNA repair hotspots. (A) Total DNA repair-associated TPMs (transcripts per kilobase million) from Repair-Seq compared with RNA-associated reads from total RNA-Seq. (B) DNA repair-associated reads from DRHs compared with RNA-associated reads from total RNA-Seq. (C) Total DNA repair-associated reads compared with RNA-associated reads from total RNA-Seq in length-normalized TADs. (D) Peak DNA repair-associated reads compared with RNA-associated reads from total RNA-Seq in length-normalized TADs. (E) Select biological process gene ontology terms for genes containing DRHs. (F) Line plot of transcription start sites (TSS) to DRHs in each gene compared with total gene length (colored by total DNA repair level).

We next looked to see if hotspots genes were significantly enriched for specific cellular processes and found that they were more correlated with genes essential for neuronal identity and function, irrespective of expression level (ex: *DLG4*, *ARC*, *GRIA4*, *GRIN2B*, *MAP2*, *HOMER1*) (Fig. 3E & fig. S7) (Table S4). Given that neural genes are typically quite long (*34*), we explored whether gene length played a role in DRH density. We compared both total repair and transcription to gene length and found that they were independent of size (fig. S8A-B). However, when we examined reads that were only from DRHs in relationship to length, we observed that the total level of repair in these sites as well as total peak density paradoxically diminished as genes grew larger (Fig. 3F & fig. S8C-D). These findings suggests that DRHs in neural genes might in part arise from the specific requirements of maintaining transcriptional elongation and splicing in genes containing large introns (*35*).

Prior reports have suggested that neuronal activity generates DSBs and the associated DNA damage marker γH2AX (phospho-histone H2A.X Ser139) in the promoters of a small subset of immediate early genes required for learning and memory to initiate transcription in mice (*12, 13*). We stimulated human ESC-iNs for 30 minutes with 50 mM KCl and allowed them 24 hrs to recover in the presence of EdU to label activity-induced DSB sites. A close examination of the promoters for activity-related genes suggested that repair occurred there under steady state with no change following stimulation and recovery (fig. S9A-B). Therefore, the lack of elevated DNA repair at these sites suggests that there might be some species-specific differences in how these genes are transcribed (*36*), that their repair might be highly reliable and not incorporate new nucleotides, or that the γH2AX that is associated with activity may not be a reliable marker of DSBs (*37*).

As cells age, the activity of DNA repair mechanisms declines (*38*), leading to an increase in genome instability in the form of information loss via somatic mutations and the accumulation of unrepaired lesions (*18, 39*). We compared the locations of somatic single-nucleotide variants (sSNVs) (*40*) identified from single neurons isolated from post-mortem humans with the DRHs we identified in ESC-iNs, and we found that they had negligible overlap (fig. S10A-B). Relative distance comparison (*41*) for DRHs showed no proximal enrichment to sSNVs (Fig. 4A & fig. S10C), suggesting mutations were occurring randomly throughout the genome, irrespective of repair efforts (*42*). We were curious as to the relative value of the genetic information that these DRHs appeared to protect. We used evolutionary conservation based on the genomic evolutionary rate profiling (GERP) score as a proxy for the relative importance of the underlying sequence (*43*). Intriguingly, DRHs often contained a single base pair under strong conservation, whereas randomly simulated peaks and sSVNs were more likely to be found at sites with negligible selective pressure (fig. S10D). We next compared the overlap of GERP-identified constrained elements (CEs) to DRHs, finding that repair was more enriched near CEs than somatic mutations (Fig. 4B & fig. S10E-F & fig. S11) (*44*). These data strongly suggest that DRHs protect essential elements from both erroneous repairs and from going potentially unrepaired.

**Fig. 4.**
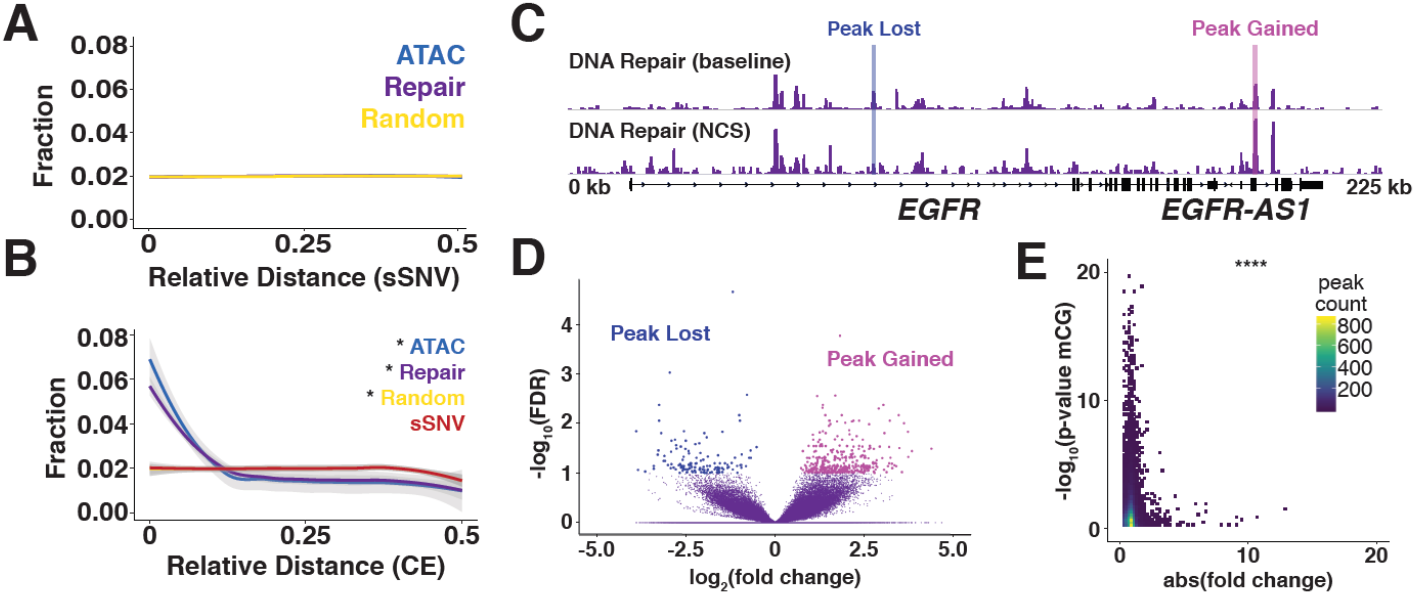
DNA repair hotspots protect evolutionarily constrained regions of the human genome from epigenetic drift. (A) Relative distance measurement from sSVNs identified from whole genome sequencing of single post-mortem human nuclei to nearest DRH or randomly placed peaks. (B) Relative distance measurement from GERP CE to nearest sSNV, DRH, ATAC-Seq peaks, or randomly placed peaks. (C) Representative browser view of DNA repair hotspots at baseline and 24 hrs after 10 min of 10 ng/mL NCS treatment demonstrates that peaks are lost and gained. (D) Volcano plot for NCS differential peaks using FDR <0.1 for DNA repair hotspots from 2 H1 and 2 H9 ESC-iNs samples. (E) Heat map of the stability (absolute fold change) of all DNA repair peaks in NCS-treated neurons compared with CG methylation changes from sorted human neurons. * p-value<0.01 by Jaccard distance test and **** p<2.47e-17 by hypergeometric test.

Aging drives fundamental changes in the epigenome - epigenetic drift - that include alternation of chromatin marks and packaging, as well as changes directly to DNA methylation patterns (*5*). Biological age is most often quantified with epigenetic clocks, i.e., changes in the methylation patterns on CG dinucleotides that are calibrated for specific cell and tissue types (*45*). Many thousand CG dinucleotides may have statistically significant methylation changes during aging; however, only a small subset of a few hundred is needed to accurately train a model for aging. Despite the accuracy of such models, no satisfying biological explanation exists as to why these DNA modifications are linked to aging (*45*). We compared the direct overlap of DRHs with CG dinucleotides from an Illumina Infinium 450K methylation array and found substantial overlap (fig. S12A). The relative distance to CG dinucleotides and CpG islands was much closer to DRHs than randomly placed peaks (fig. S12B-C). Using CG dinucleotides that exhibited methylation changes statistically associated with aging neurons from human prefrontal cortex (*46*), we found some direct overlap with DRHs and a closer relative distance than random (fig. S12D-E).

Genome instability in the form of DSBs is thought to be a primary driver of biological aging (*47*). We treated our neurons with the radiomimetic DNA-damaging agent neocarzinostatin (NCS) to assay the changes to DRHs following injury. Acute NCS treatment triggered both the gain and loss of DRHs in neurons in a largely stochastic fashion, though at the dosage used relatively few peaks demonstrated consistent change (Fig. 4C-D & fig. S12F) (Table S5). In the context of aging, genome instability would potentially redistribute repair efforts away from hotspots to other locations in the genome, similar to what we observed with NCS treatment (*48*). We compared absolute fold change for NCS-treated samples with statistically significant CG methylation sites and found that the most stable sites were those most likely to be associated with the epigenetic clock (Fig. 4E & fig. S12G). Therefore, as DNA repair capacity declines with age, many of these sites might become less maintained as pathways become overtaxed, and subsequently more susceptible to changes in methylation status.

Collectively, incorporation of the click nucleoside analog EdU into the genome by repair polymerases has provided a useful tool to visualize the locations of DNA repair in neurons as well as a means to isolate genome fragments and sequence their locations. Our results conclusively demonstrate the existence of recurrent DRHs in post-mitotic neurons and suggested that they played a key role in neuron identity and function. Going forward, Repair-Seq will be a powerful tool to explore how age and disease can disrupt genome integrity in the nervous system. Finally, whether DRHs are a unique feature of neuron genome protection, specific developmental lineages, non-dividing cells, or are limited to only some long-lived species remains and open question. The possible discovery of these sites in other cell types might further aid in our understanding of how age-related changes in their organization could drive differential aging or the development of disease in other tissue types.

## Acknowledgments

The authors would like to thank L. Moore, I. Guimont, and K.E. Diffenderfer, W. Travis Berggren for technical assistance and M.L. Gage for editorial comments. They would also like to acknowledge the Salk Institute Stem Cell Core, Waitt Biophotonics Core, and Next Generation Sequencing Core for technical support.

## Author contributions

D.A.R. conceived of the project, generated the data, helped analyzed the results, and supervised the project in coordination with N.H., C.K.G., and F.H.G. Repair- and RNA-Seq libraries were generated by D.A.R., G.C., C.A.M., J.H.O. ATAC- and ChIP-Seq experiments were performed by J.C.M.S, and Hi-C experiments were performed by S.C. Additional experiments and reagents were contributed to the study by J.R.J., A.S.R., E.C.T., S.L., S.T.S. Analysis of Repair-, ATAC-, ChIP- and RNA-Seq was performed by P.J.R. and S.B.L. Analysis of Hi-C was performed by P.J.R. and J.R.D. DNA methylation aging analysis was performed by A.T.L and S.H. The manuscript was written by D.A.R., P.J.R., and F.H.G, and edited by J.C.M.S., J.R.J., J.R.D., and C.K.G.

## Data and materials availability

Primary data will be made available on GEO upon publication and is presently available by request.

## Supplementary Materials

## Figures S1-S12

**fig. S1.**
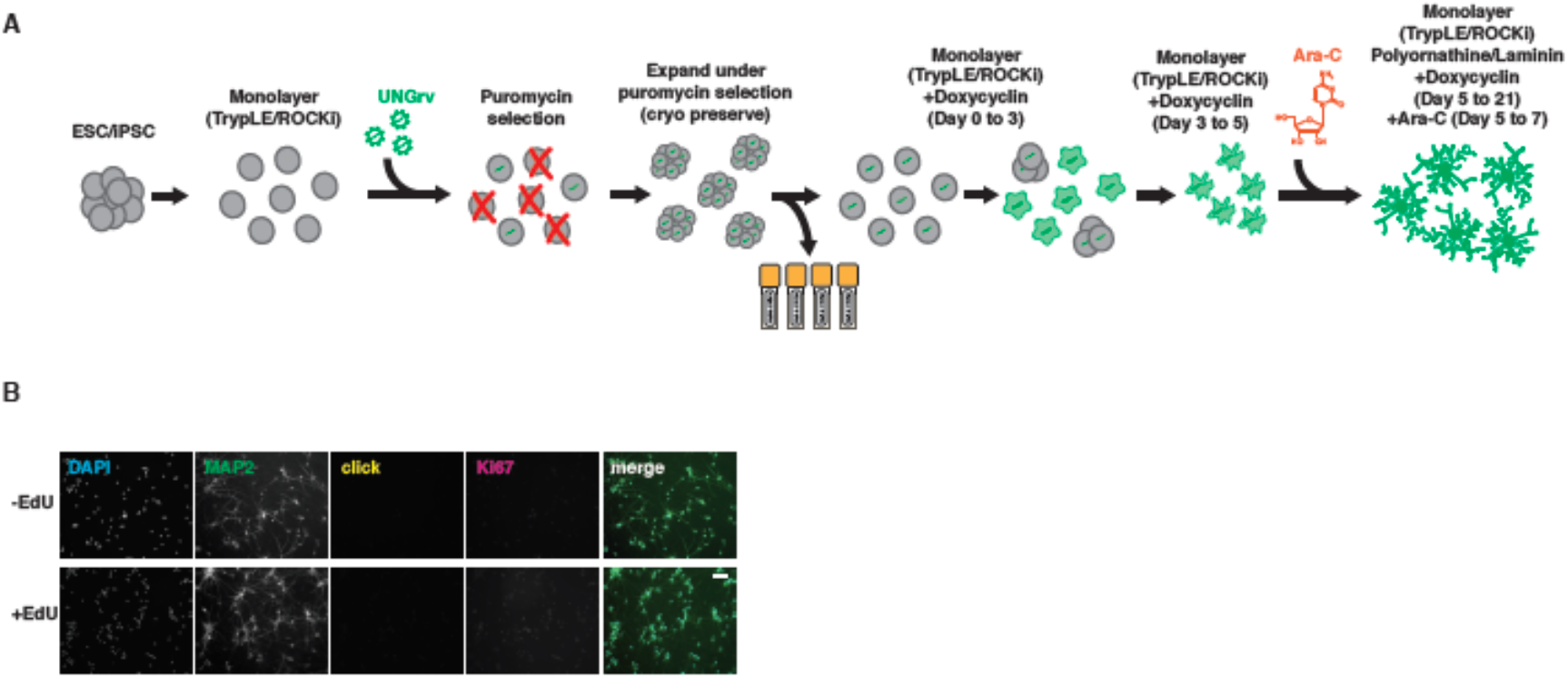
Schematic for assembly and sequencing of Repair-Seq libraries. (A) Schematic for the production of pure ESC-iNs without flow sorting. (B) Representative images of ESC-iNs demonstrate no EdU positive nuclei (fed 24 hrs) and low levels of Ki67 staining (<5%; background) indicate that all cells are non-dividing during this period. Scale bar is 10 microns.

**fig. S2.**
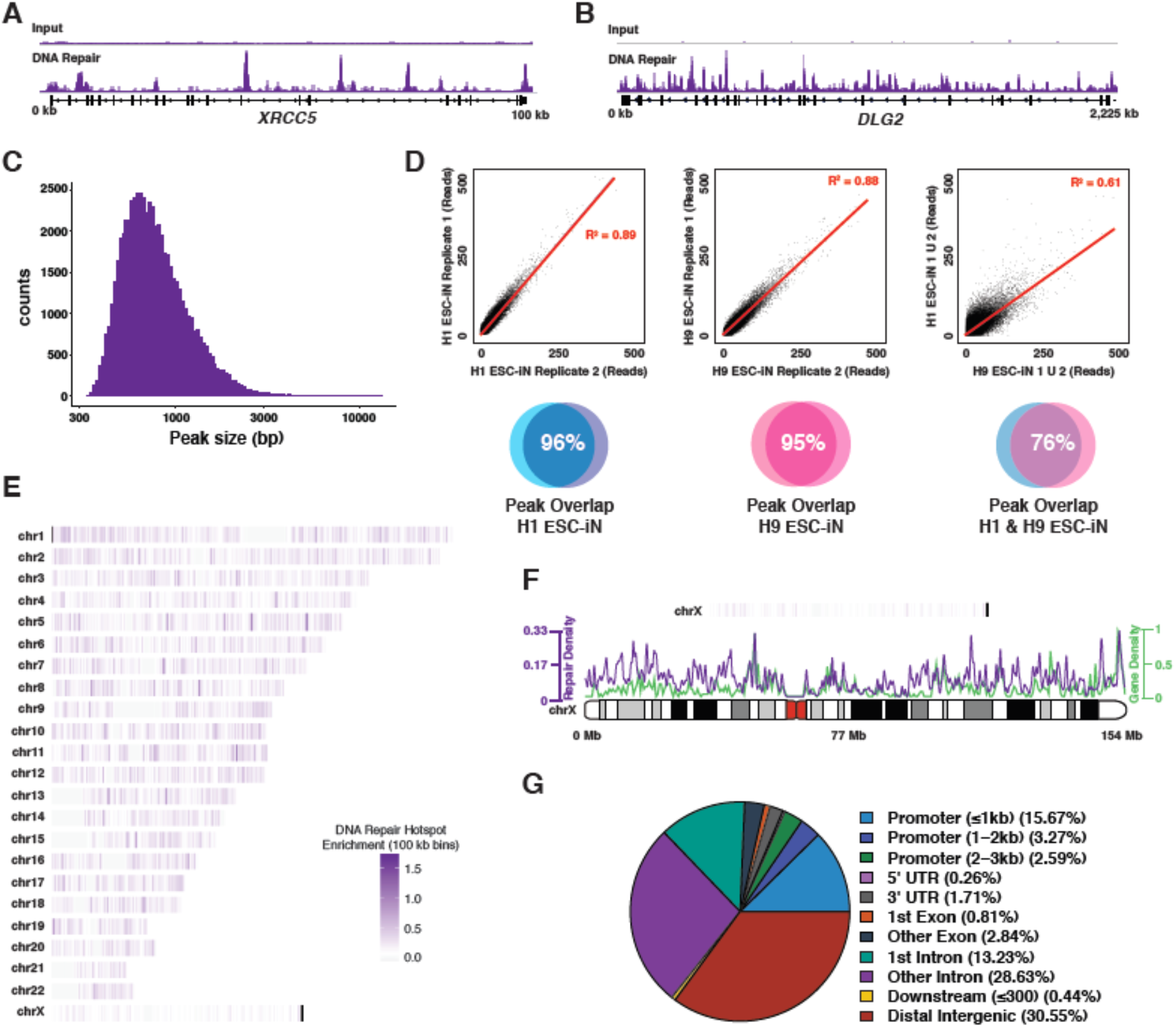
Genomics of stable genome repair hotspots in post-mitotic human neurons. (A-B) Further examples of DNA repair hotspots in the *XRCC5* and *DLG2* loci. (C) Histogram of genome repair hotspot peak widths. (D) Reproducibility of Repair-Seq peaks in normalized read counts for biological replicates of H1 and H9 ESC-iNs. (E) Genome map of DNA repair hotspots in ESC-iNs. (F) Detailed view of DNA repair hotspots (purple) and gene density (green) on Chromosome X. (G) Genome annotations for DNA repair hotspots show distributions primarily in promoters, gene bodies, and intergenic regions.

**fig. S3.**
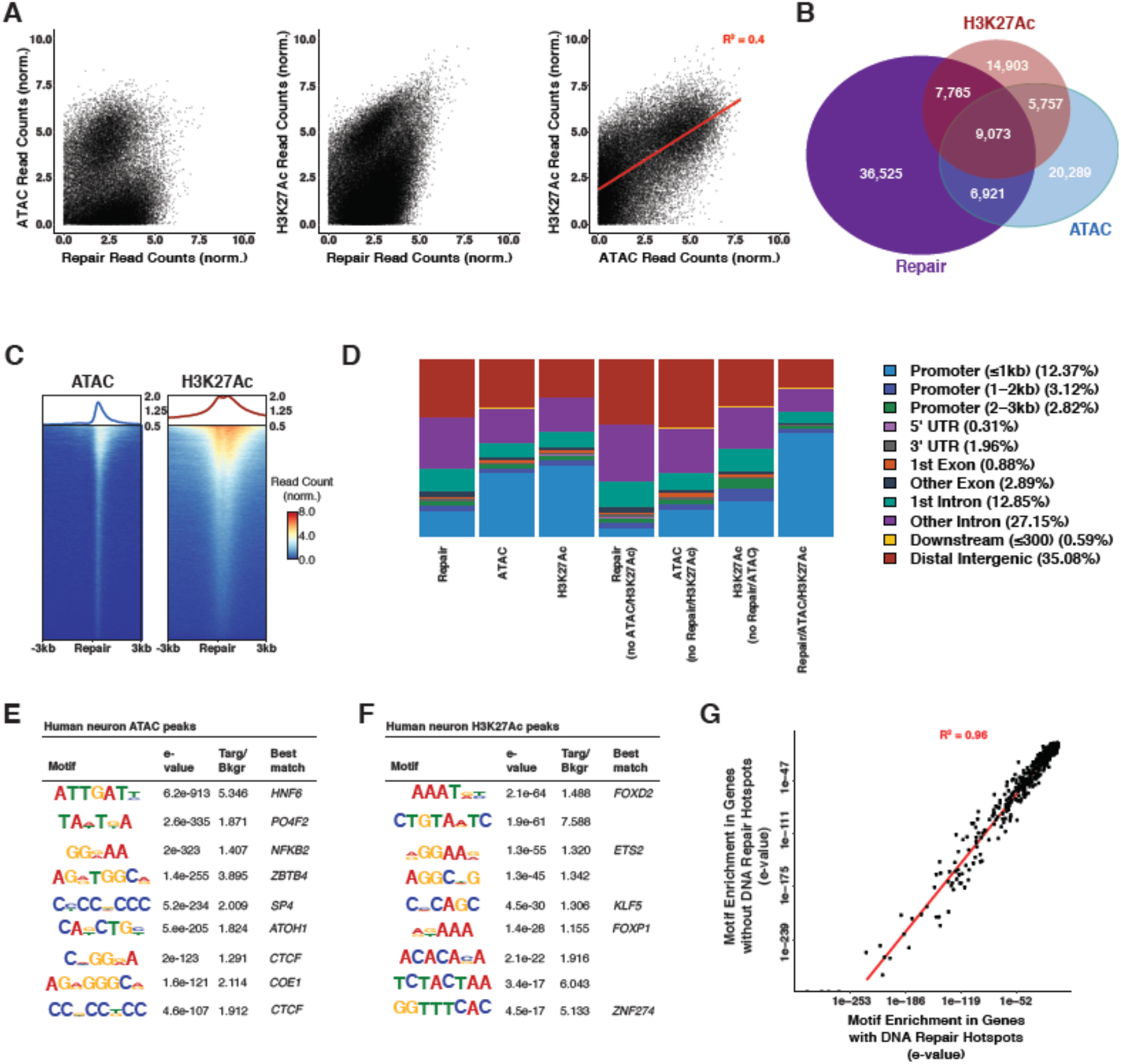
Chromatin accessibility is a primary driver of DNA repair in neurons. (A) Scatter plots of Repair-Seq vs ATAC-Seq, Repair-Seq vs H3K27Ac ChIP-Seq and ATAC-Seq vs H3K27Ac ChIP-Seq. (B) Repair-Seq, ATAC-Seq, and H3K27AC ChIP-Seq peak overlaps. (C) TSS plots for ATAC-Seq and H3K27Ac ChIP-Seq peaks centered on nearest Repair-Seq peak. (D) Genomic annotations for Repair-Seq, ATAC-Seq, H3K27Ac ChIP-Seq peak overlaps. (E) DNA sequence motifs identified *de novo* in Repair-Seq peaks using all ATAC-Seq peaks as background. (F) DNA sequence motifs identified *de novo* in Repair-Seq peaks using all H3K27Ac ChIP-Seq peaks as background. (G) Comparison of the enrichment *de novo* DNA sequence motifs identified in DNA repair hotspots in genes that have hotspots and lack hotspots.

**fig. S4.**
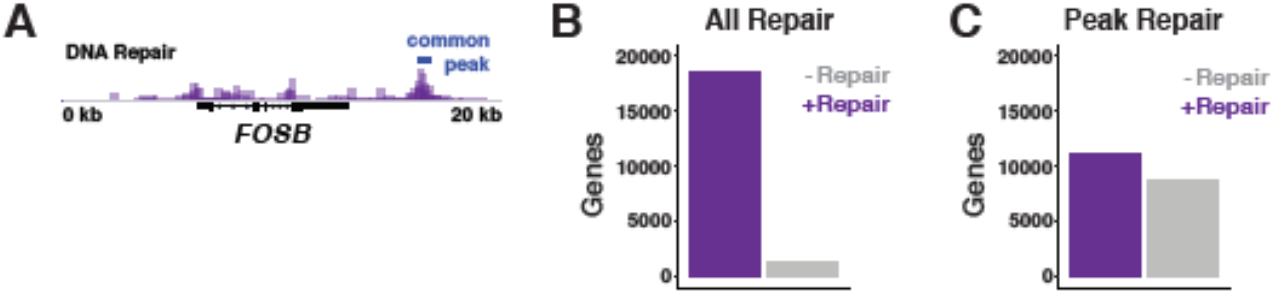
Quantification of genes with DNA repair. (A) DNA repair in *FOSB* locus. (B) Bar plot displaying the number of protein-coding genes that have DNA repair-associated reads. (C) Bar plot displaying the number of protein-coding genes that have DNA repair-associated reads found in DNA repair hotspots.

**fig. S5.**
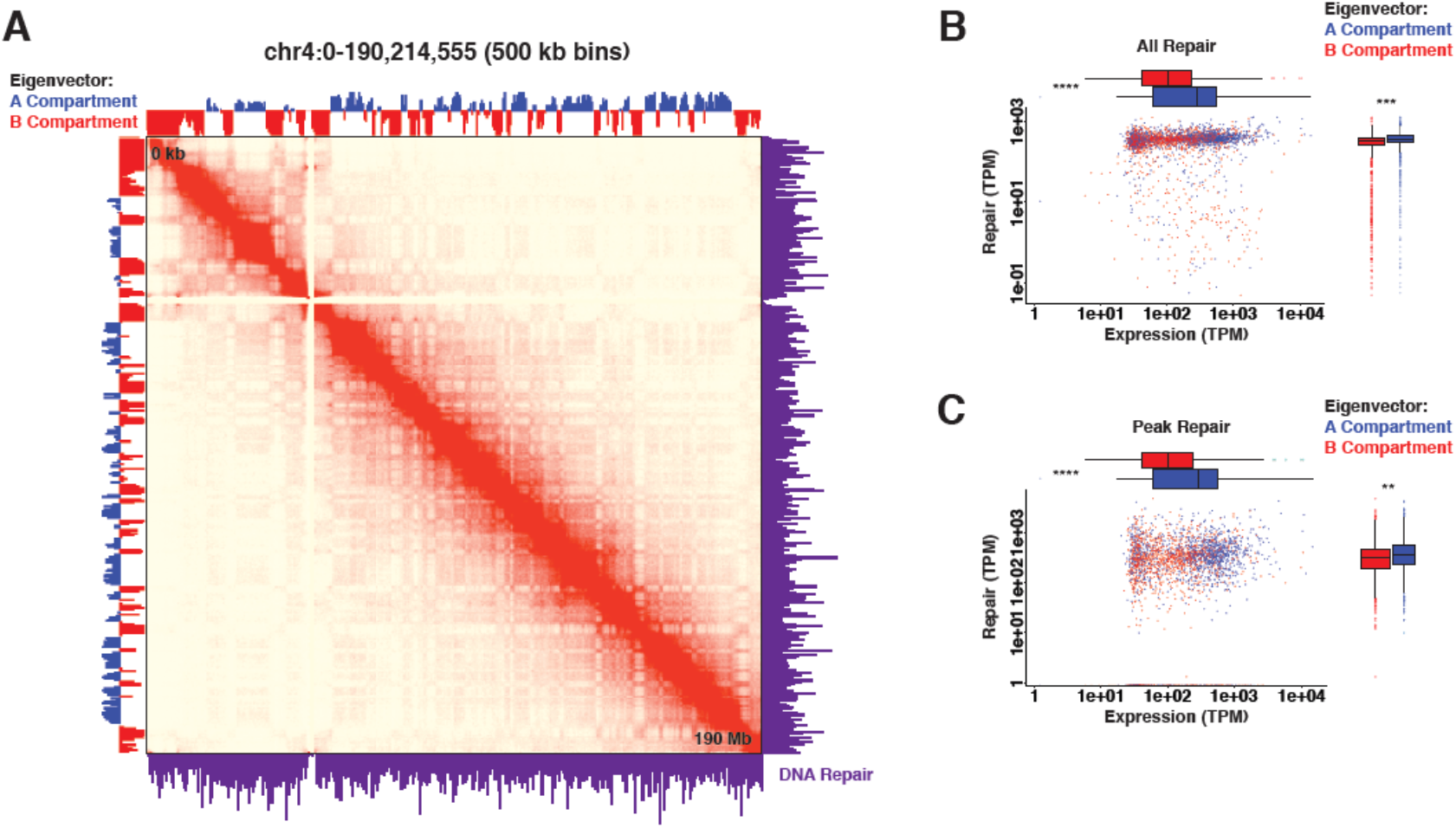
Hi-C A/B compartments are enriched for DNA repair. (A) Representative Hi-C data for chromosome 4 displaying eigenvalues corresponding to A/B compartments and DNA repair from Repair-Seq. (B) Box and scatter plots of all DNA repair associated reads compared with transcription in Hi-C A/B compartments. (C) Box and scatter plots of all DNA repair peak associated reads compared with transcription in Hi-C A/B compartments. **** p-value<2.8e-44, *** p-value<8.5e-19, ** p-value<3.8e-13 by Wilcoxon test.

**fig. S6.**
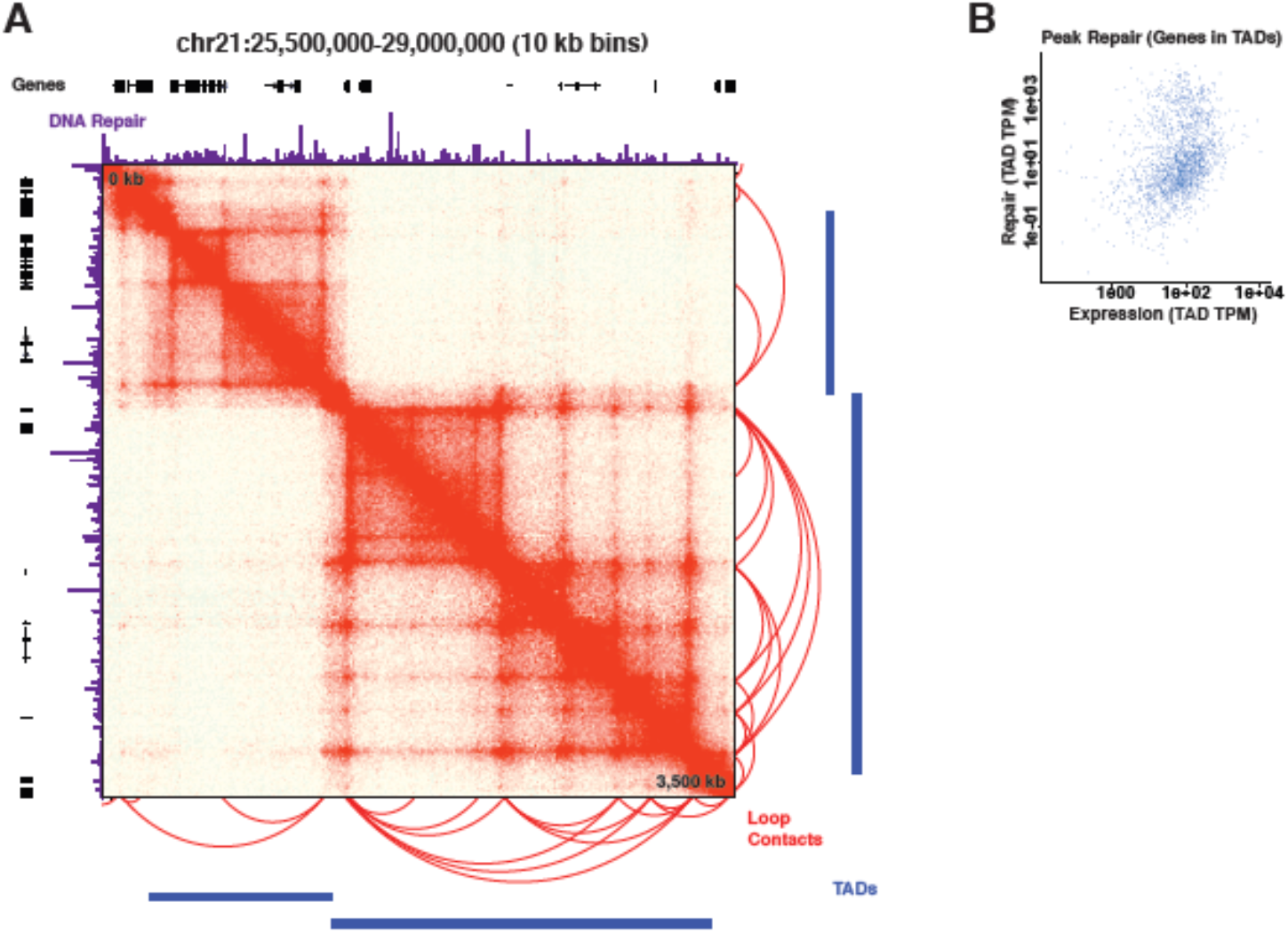
TADs are not preferentially enriched for DNA repair. (A) Representative Hi-C data from chromosome 21 displaying loop contacts, TADs, DNA repair and genes. (B) Scatter plot of DNA repair-associated reads found in SGRHs in genes compared with total expression normalized to TAD width.

**fig. S7.**
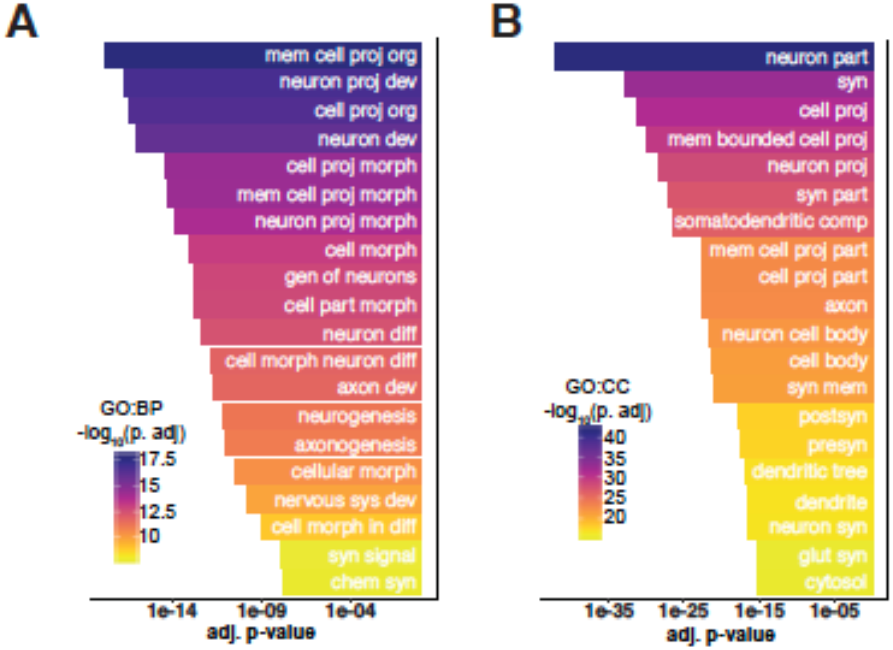
Complete GO terms for biological process and cellular component for genes with DNA repair hotspots. (A) Top 20 biological process GO terms for genes with SGRHs. (B) Top 20 cellular component GO terms for genes with SGRHs.

**fig. S8.**
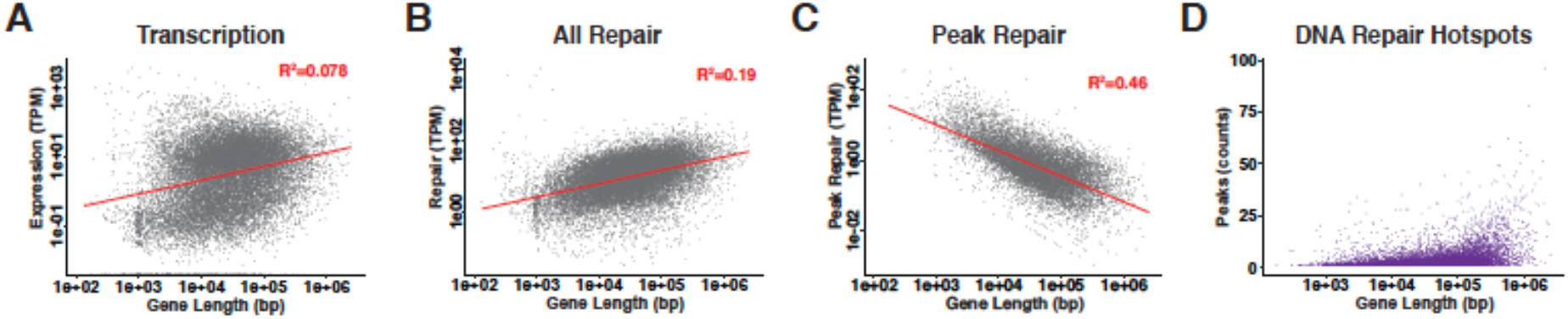
Length dependency of DNA repair hotspots. (A) Normalized transcription compared with gene length. (B) Normalized DNA repair compared with gene length. (C) Normalized DNA repair in peaks compared with gene length. (D) Number of DNA repair hotspots compared with gene length.

**fig. S9.**
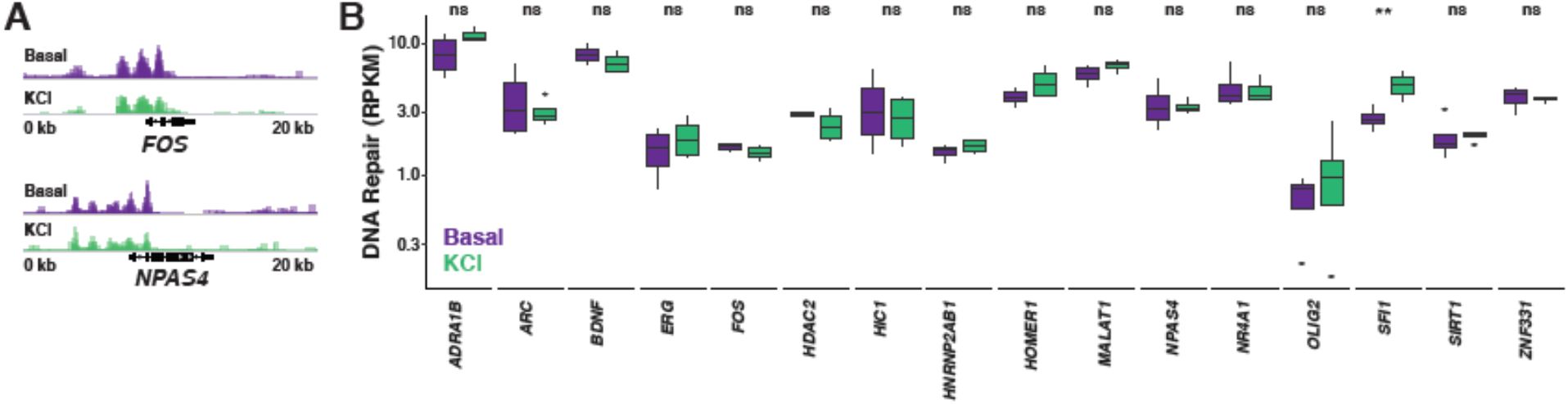
Neuronal depolarization does not increase DNA repair levels in the promoters of immediate early genes. (A) Promoter region of *FOS* and *NPAS4* loci in basal conditions (purple) and with the addition of 50 mM KCl for 30 minutes and 24 hr of recovery (green). (B) TPM box plots for the promoters of genes thought to have activity-induced DSBs found in mouse cortical neuron culture (*FOS*, *MALAT1*, *NPAS4*, *ERC*, *OLIG2*, *NR4A1*, *HOMER1*, *NR4A3*, *HDAC2*, *HNRNP2AB1*, *SIRT1*) and human-specific activity-related genes (*BDNF*, *ARC*, *HIC1*, *LINC00473*, *ZNF331*, *ADRA1B*). ** p-value<0.01 by Wilcoxon test.

**fig. S10.**
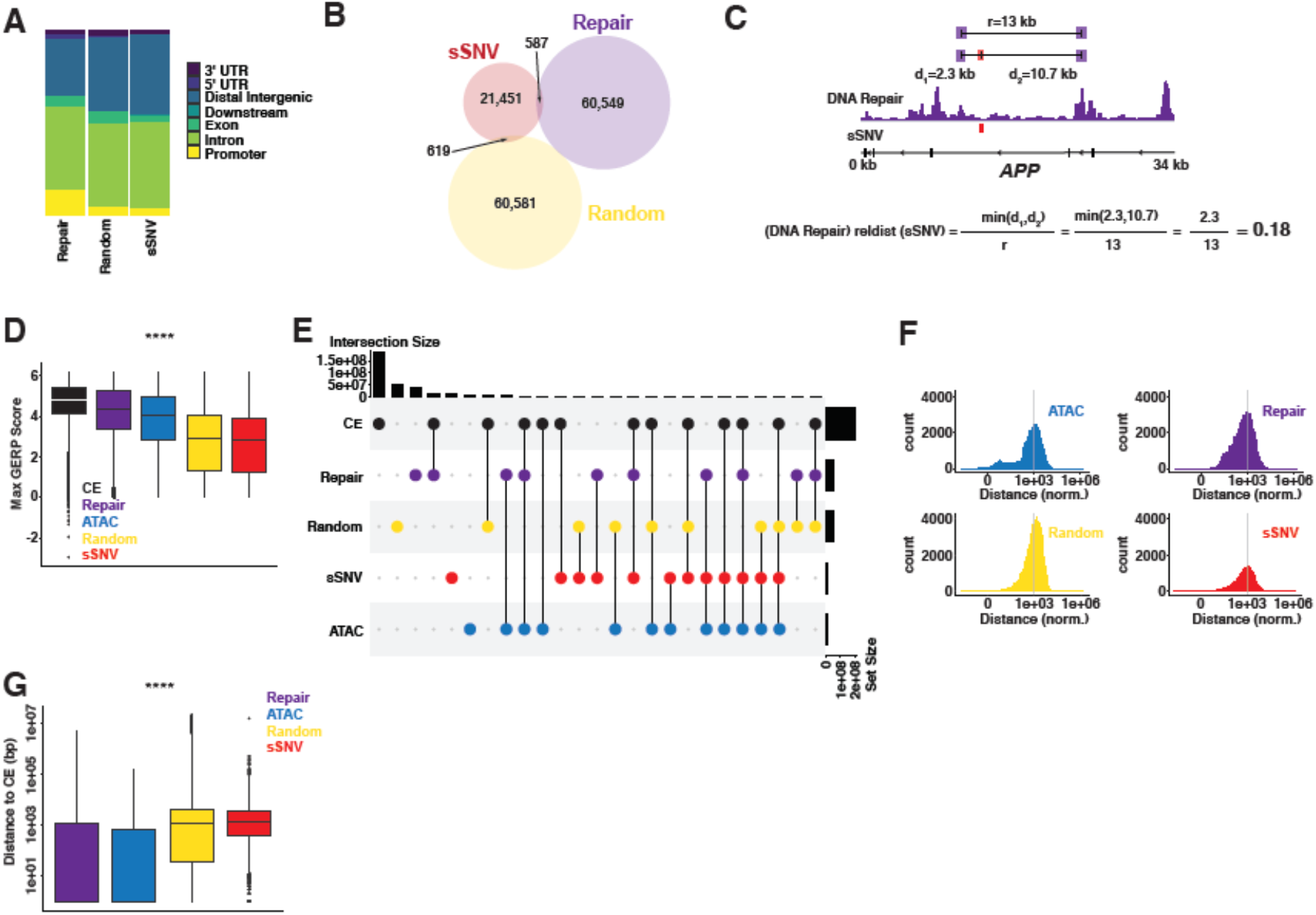
DNA repair hotspots correlate with key genomic regions. (A) Genomic distributions of DNA repair hotspots, random peaks, and sSNVs from human neurons. (B) Venn diagram for overlaps of Repair-Seq peaks with sSNVs and Random peaks with sSNVs. (C) Schematic for relative distance (reldist) function from bedtools. (D) Max GERP score for CEs, Repair-Seq peaks, ATAC-Seq peaks, Random peaks, and sSNVs. (E) Upset plot of intersections for CEs, Repair-Seq peaks, Random peaks, ATAC-Seq peaks, and sSNVs. (F) Interpeak distances for ATAC-Seq peaks, Repair-Seq peaks, Random peaks, and sSNVs normalized (bp*peaks/1e6). (G) Box plots for absolute distances for CE to Repair-Seq peaks, CE to ATAC-Seq peaks, CE to Random peaks, and CE to sSNVs. **** p-value<2e-16 by one-way ANOVA.

**fig. S11.**
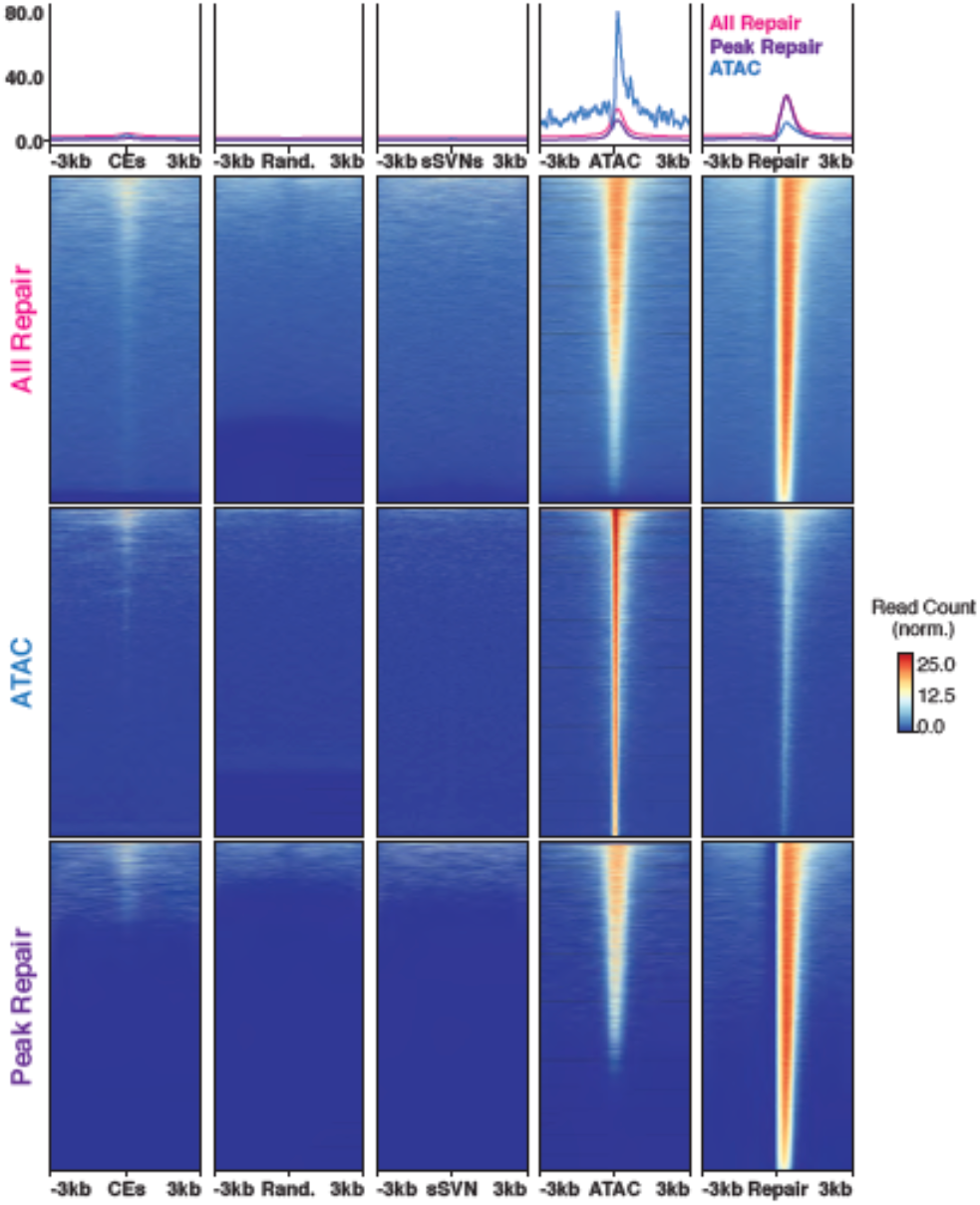
Heatmaps for All Repair, Peak Repair, and ATAC-Seq. Heatmaps centered on CEs, Random peaks, sSNVs, ATAC-Seq peaks, and Repair-Seq peaks compared with all Repair-Seq reads, ATAC-Seq peaks, and Repair-Seq peaks.

**fig. S12.**
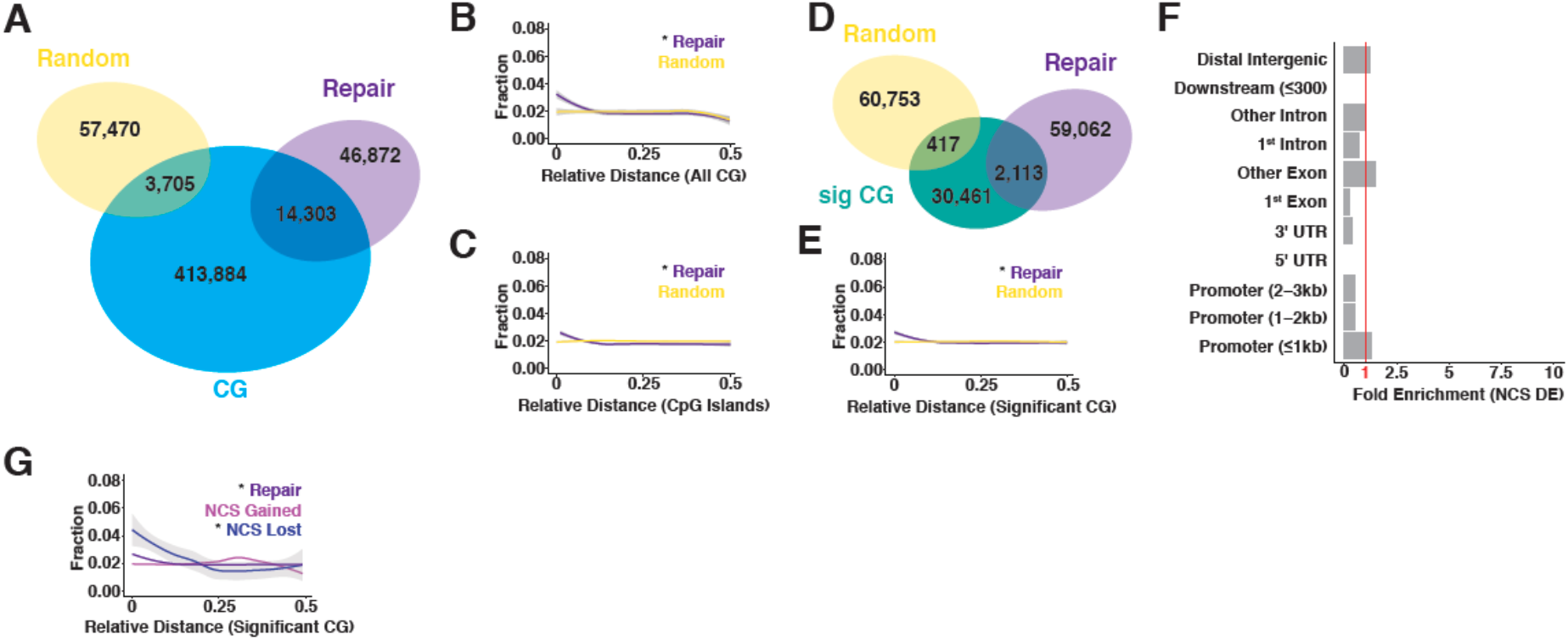
DNA damage and epigenetic drift. (A) Overlaps between CG dinucleotides on Illumina Infinium 450K methylation array, Repair-Seq peaks, and Random peaks. (B-C) Relative distance plot of Repair-Seq and Random peaks to CG dinucleotides from an Illumina Infinium 450K methylation array and CpG islands in the human genome. (D) Overlaps between CG dinucleotides that are significantly associated with methylation changes in aging human neurons and Repair-Seq or Random peaks. (E) Relative distance from significant CG dinucleotides to either Repair-Seq or Random peaks. (F) NCS peaks that are gained and lost largely at random when normalized to existing DRHs. (G) Relative distance measurement from NCS gained and lost sites to significant CG dinucleotides. * p-value<0.01 by Jaccard distance test.

